# Fighting Sleep: Minute-by-Minute Driver Fatigue from a Dry-Electrode EEG Headband

**DOI:** 10.64898/2026.11.18.767417

**Authors:** Ryan S. Bohluli, Avaneesh Babu, Lucas R. Wein, William M. Blake

## Abstract

Driver-facing cameras often catch fatigue too late. We asked whether a four-electrode dry headband, which picks up eye movements electrically and head tilt with a motion sensor, can follow fatigue minute by minute. Nine participants wore the headband for twelve sessions of a 50-minute monotonous task at a driving simulator station, without driving, answering one cue a minute and rating their sleepiness every five minutes, with a webcam on the face for comparison. An index built without training from blink length, slow eye movements and head tilt, read against each participant’s own first eight minutes, followed fatigue beyond elapsed time across the nine participants (0.32, p = 0.002), above zero in eight of them and at zero in the ninth. With its alarm line set for a 10 percent false-positive rate on not-sleepy minutes, it caught 91 percent of the minutes rated as fighting sleep, where the standard camera eye-closure measure (PERCLOS, the share of each minute with the eyes at least 80 percent closed), set the same way on the same minutes, caught 33 percent (67 minutes from four participants). Averaged over the six recordings that reached fighting sleep, the index crossed its alarm line about three minutes before the first minute rated as fighting sleep; in the five sessions with video its alarm had sounded by then in four, where the camera’s sounded 6 to 13 minutes after in four and never in the fifth. The index and its alarm were then fixed and taken into a car, on one outing by one participant. Through a 33-minute rested drive the headband returned a value for every minute and raised no alarm. With the car parked and the driver doing the laboratory task, its alarm sounded nine minutes before he fell asleep, as seen on video; the camera’s, set the same way, came eight minutes later, with one minute to spare. Signal amplitude at the electrodes was no higher in the parked car than in the laboratory, and with the car moving the electrodes that carry the index stayed in range 99 percent of the time.

## I. Introduction

**D**RIVER fatigue kills, and the official counts understate how often. Police reports in the United States attributed about 90,000 crashes and 824 deaths to drowsy driving in 2015, and the agency that compiles them notes that drowsiness is underreported [1]. When drivers were filmed in their own cars, eye closure in the minutes before a crash put drowsiness in 8.8 to 9.5 percent of all crashes [2]. A later estimate put a drowsy driver in 17.6 percent of fatal crashes involving passenger vehicles from 2017 to 2021, an estimated 29,834 deaths in five years [3]. The risk climbs steeply with lost sleep. Drivers who had slept less than four hours in the past day crashed at 11.5 times the rate of those who had slept seven or more, a risk comparable to a blood alcohol concentration of 0.12 to 0.15, against a legal limit of 0.08 [4]. The cost of fatigue-related fatal and injury crashes has been put at 109 billion dollars a year, before property damage [5].

Crashes involving large trucks killed 5,936 people in the United States in 2022 and injured an estimated 160,608, and seven in ten of the dead were in other vehicles [6]. Trucker fatigue is part of that toll. In the national study of large-truck crashes with a death or injury, fatigue was recorded for 13 percent of the trucks [7], [8]. Where the critical reason for the crash lay with the truck, the driver had fallen asleep or was otherwise physically impaired in 12 percent of cases, and had failed to recognise the situation, as through inattention or distraction, in a further 28 percent [8]. The rule that guards against trucker fatigue counts hours, allowing 11 hours of driving inside a 14-hour window after 10 hours off duty [9]. It does not measure the state of the driver.

Drivers are a weak safeguard against their own fatigue. In a three-hour simulated night drive by 90 people, three quarters of those who rated their drowsiness low were moderately or highly drowsy by eye closure, and of those who rated themselves extremely drowsy, more than three quarters drove on past a rest area [10].

The response in vehicles has been the driver-facing camera. It tracks the eyelids, and the standard measure taken from it is PERCLOS, the share of time the eyes are mostly closed [11]. Camera systems built on it have been described for two decades [12], [13], and assessment programmes now score cars on them, requiring a driver to be classed as drowsy by a rating of 8 on the nine-point Karolinska scale [14]. A camera reads the eyelids and head optically; a headband can pick up eye movements electrically and head tilt with a motion sensor.

In the laboratory, sleepiness is read from the body. The EEG and the electrical signal of the eyes change as a person fights sleep [15], [16], and blinks lengthen [17], [18]. These signs follow the rating a person gives of their own sleepiness and their performance [19], [20], and they have been recorded on the real road [21], [22]. Carrying them into a vehicle means electrodes on the head. Road studies have worn headsets of wetted electrodes [23], [24], and work on smaller montages is recent [25].

This paper asks a narrower question than whether EEG can detect drowsiness. Can a dry headband of four electrodes follow fatigue minute by minute, and does it tell more than a clock or a camera would? The clock matters because in a single unbroken session fatigue rises with time [21], [26], so any signal that drifts will appear to follow it. The camera matters because it is the instrument already in vehicles. Reported accuracy in this field also depends on how a detector is validated [27], [28], so the index tested here has no weights learned from the data.

Nine participants wore such a headband for twelve sessions of a 50-minute monotonous task at a driving simulator station, with a webcam on the face throughout. This paper makes four contributions. First, an index built from blink length, slow eye movements and head tilt, with no training and no clock, follows fatigue beyond elapsed time across the nine participants, above zero in eight of them and at zero in the ninth. Second, with its alarm line set for a 10 percent false-positive rate on not-sleepy minutes, the index catches 91 percent of the minutes participants rated as fighting sleep, where the standard eye-closure measure from the camera, set the same way on the same minutes, catches 33 percent. Third, the signal comes from the eye and head measures; EEG band power from the same electrodes did not improve the index on these recordings. Fourth, the index and its alarm line were fixed and written down before any recording in a car, and on one outing by one participant the alarm stayed quiet through a rested drive and sounded nine minutes before the driver, by then parked and doing the laboratory task, fell asleep, and eight minutes before the camera’s alarm did. Signal amplitude at the electrodes was no higher in the parked car than in the laboratory (Sections V-G and VI-F).

Section II reviews prior work, Section III describes the headband, Sections IV and V the study and its analysis, Section VI the results, and Section VII what they do and do not support.

## II. Background and Related Work

This section covers two bodies of prior work: EEG recorded from a few electrodes at the side and front of the head, and the recording of EEG and sleepiness in moving vehicles. Table I places the present study among them by number of electrodes and by setting.

**TABLE I.** EEG studies of driver state, by electrode count.

| **Study** | **Meas. el.** | **Interface** | **Site and setting** | **Label rule** | **Unit of analysis and split** |
|---|---|---|---|---|---|
| Wang 2025 [24] | 1 | Semi-dry, conductive liquid | O1 scalp; on road, driving | KSS | Epoch; split not stated |
| Yao 2025 [37] | 1 | Semi-dry, conductive liquid | O1 scalp; on road, driving | KSS 6 or below against above 7 | 2 s sample; shuffled 80/20, then 5-fold |
| Zhang 2026 [38] | 2 | Semi-dry, conductive liquid | O1 and O2 scalp; on road, 3 h drive | KSS 7 or above | 8 s epoch; six folds, two participants held out in each (80/20 also stated) |
| Minhas 2025 [39] | 6 | Not stated | Six scalp sites incl. F4 and O2; simulator | Visual scoring of facial video | Epoch; subject-specific thresholds |
| Zhu 2020 [40] | 8 | Dry | Eight scalp sites; simulated driving | Recording condition, awake or drowsy | 1 s sample; shuffled, random split |
| Schwendeman 2022 [41] | 10 | Dry | In-ear canal and concha; indoors, at a laptop | RT above 2.5 s, alert unless a drowsy Likert item accompanied it | Event, 3 consecutive min; leave-one-user-out |
| Kaveh 2024 [25] | 11 | Dry, wet mastoid ground | In-ear canal and concha; lab desk | RT above twice own first 5 min AND Likert above 5 of 10 | Event; leave-one-trial-out; rule not stated |
| Di Flumeri 2018 [42] | 12 | Not stated | Scalp; on road, urban | Cognitive workload, not fatigue | Workload neurometric |
| van Klaren 2024 [43] | 18 | Gelled | Around-ear, cEEGrid; lab, sleep restriction | KSS 1 to 9 and a 10 min PVT | No classifier; repeated-measures ANOVA, rmcorr |
| Liu 2023 [44] | 21 | Not stated | Scalp; on road, car, six-lane city avenue | Cognitive workload, not fatigue | ERSP and band power |
| Simon 2011 [45] | 43 | Wet | Scalp, nose ref; on road, 430 to 480 km highway | Drive aborted from fatigue; first vs last 20 min | 1 s segments, 750 ms overlap; ANOVA |
| This work | 4 | Dry | Headband, TP9, TP10, AF7 and AF8; simulator station | Rank index of RT and KSS; KSS 9 at the strictest | 1 min; fixed index, folds by participant |

*Measurement electrodes exclude the reference, the ground, and any driven electrode.*

### A. What has been recorded, and with what

Recording EEG from a few electrodes at the side of the head is established practice. Electrodes in and around the ear, close to where the temporal electrodes of a headband sit, resolve auditory steady-state responses, the P300 and the attenuation of the alpha rhythm on eye opening [29], [30]. Dry electrodes in the ear have recorded whole nights of sleep at home [31]. In-ear conductive-polymer electrodes have also recorded the electrocardiogram, in rats, with heart rate close to that from chest electrodes [32].

A few electrodes at the side and front of the head see less than a full scalp montage. The usual check that a montage records cortex and not noise is the ratio of alpha band power with the eyes closed to that with the eyes open. The nearest published figures come from electrodes in and around the ear, close to the temporal sites of a headband: about 1.23 to 1.25 between the two ear canals (differences of 0.89 and 0.96 dB) [33], 1.51 to 2.44 for canal electrodes referenced to the concha [34] and 2.48 around the ear [35], against a reported mean of 1.5 and a range from 0.8 to 3.0 on occipital scalp [19]. Between 10 and 25 percent of participants show no usable increase at the ear at all [36].

Table I sets out the studies that record EEG from a person operating or attending to a driving task, ordered by the number of measurement electrodes. Two properties organise it: how much hardware is in contact with the head, and the setting in which the recording took place. On the road, prior work sits at the two extremes, with full wet caps and with one or two occipital electrodes carrying a conductive liquid. The ear montages, at ten to eighteen electrodes, were recorded at a desk or in the laboratory. This work records four dry electrodes at a simulator station.

### B. Recording in operational vehicles

Sleepiness has been recorded physiologically on the real road for more than a decade. Sandberg et al. recorded EEG, eye movements and blink duration from 18 drivers over 55 km of rural highway by day and at night, with a sleepiness rating every five minutes [21]. Hallvig et al. compared real driving with a simulator that reproduced the same road, and found sleepiness higher in the simulator [22]. Safety in such studies rests on the vehicle and the route. One motorway study used a car with dual controls and a test leader in the front passenger seat [46], and a recent study with a wearable headset of eight water-based electrodes drove on sites without traffic [23]. Reduced montages have been tried in driving as well, down to a single semi-dry occipital channel [24]. No study was found that recorded a dry headband of a few electrodes, together with an index built from eye and head signals, in a moving car.

## III. nstrument

The recordings were made with a four-electrode dry headband.

### A. Montage and electrodes

The headband has two temporal electrodes, at TP9 and TP10, and two forehead electrodes, at AF7 and AF8, all referenced to Fpz (Fig. 1). The forehead pair supplies the eye signs of Section V-A and the temporal pair the EEG sign. Recordings from a few electrodes in and around the ear, close to the temporal sites, resolve the alpha rhythm and evoked responses [29], [30].

**Fig. 1.**
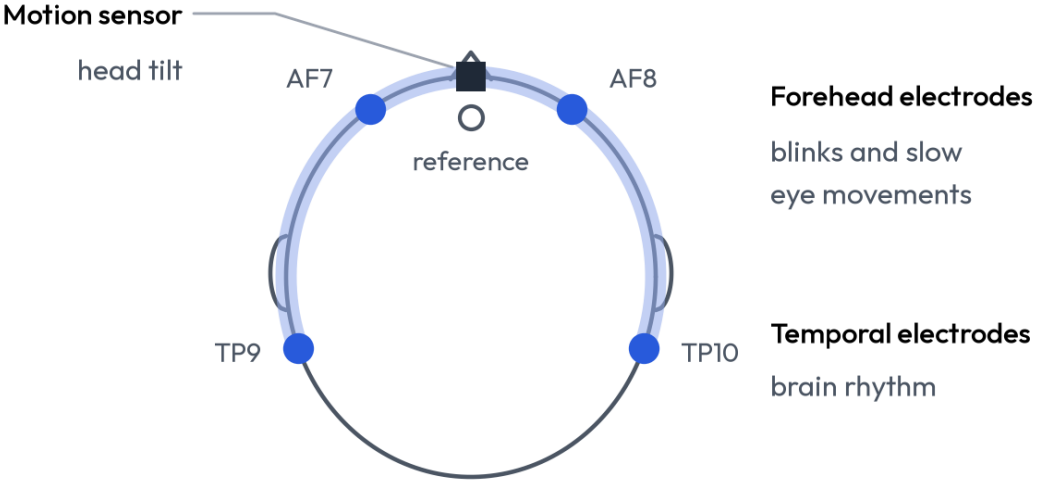
Sensor positions. The headband seen from above. Two forehead electrodes (AF7, AF8) pick up blinks and slow eye movements, two temporal electrodes (TP9, TP10) record brain rhythm, and a motion sensor at the front of the band measures head tilt. All electrodes share a reference at the centre of the forehead. The headset index is computed from the forehead electrodes and the motion sensor.

### B. Acquisition

EEG is sampled at 256 Hz, and the accelerometer and gyroscope of the headband at 52 Hz. Two units of the same model were used, with the same acquisition settings. Electrode contact was judged from the rating that the acquisition software gives each electrode, and no impedance measurement was made. The streams of the headband are stamped on arrival at the bench computer, and task events are stamped by the simulator.

## IV. Study Design

### A. Design overview

The study adapted the drowsiness task of Kaveh et al. to a four-electrode dry headband at a driving simulator station, with a webcam on the participant’s face throughout as a camera reference. Nine participants contributed twelve laboratory recordings between 3 and 15 November 2026 (Table II).

**TABLE II.** Recording settings.

| **Setting** | **Setting** | **Task** | **n** |
|---|---|---|---|
| Laboratory | Driving simulator station, seated | Reaction-time task | 9 people, 12 recordings |
| Driving | Personal passenger car: driven on local streets, then parked outside with the engine idling | Rested drive, then the reaction-time task | 1 person, 2 recordings |

Each cue is one of four directions, answered by pressing the matching one of four buttons. Sleepiness is rated on the standard nine-point Karolinska Sleepiness Scale. The main block is a fixed 50 cues over 50 minutes. Sessions take place at a driving simulator station, and a member of the study team stays in the room, seated behind the participant and silent. An eyes-open and eyes-closed block is recorded before and after the task. Three participants returned for a second session.

The analysis reported here was developed on the data while collection was under way. After data collection the headset index of Section V-B, the top-of-scale definition of fatigue and the alarm line were frozen, and they are scored unchanged on the driving trial (Section IV-F). The rule was written down and committed to the study’s repository on 15 November 2026, after the last laboratory recording and two days before the driving trial.

### B. Participants

Nine adults took part, aged 21 to 24, seven men and two women. Three of them returned for a second session; following Kaveh et al., repeat sessions were at least 24 hours apart. That gives twelve recordings. A repeat session does not add to the participant count: wherever a statistic is a mean over people, a participant’s sessions are averaged first, and every cross-validation fold is formed by participant, so a participant’s data never sits on both sides of a split [27].

Sessions began between 9 a.m. and 5 p.m. Participants were asked to have a normal night’s rest and to avoid caffeine and exercise beforehand, and each confirmed on arrival that they had.

### C. Session procedure

Sessions were run at a driving simulator station (Fig. 2), but participants did not drive: the screen was blank between cues. After intake the headband was fitted and its contact checked with the acquisition software’s rating of each electrode. The fit was adjusted until every electrode was rated good before recording began.

**Fig. 2.**
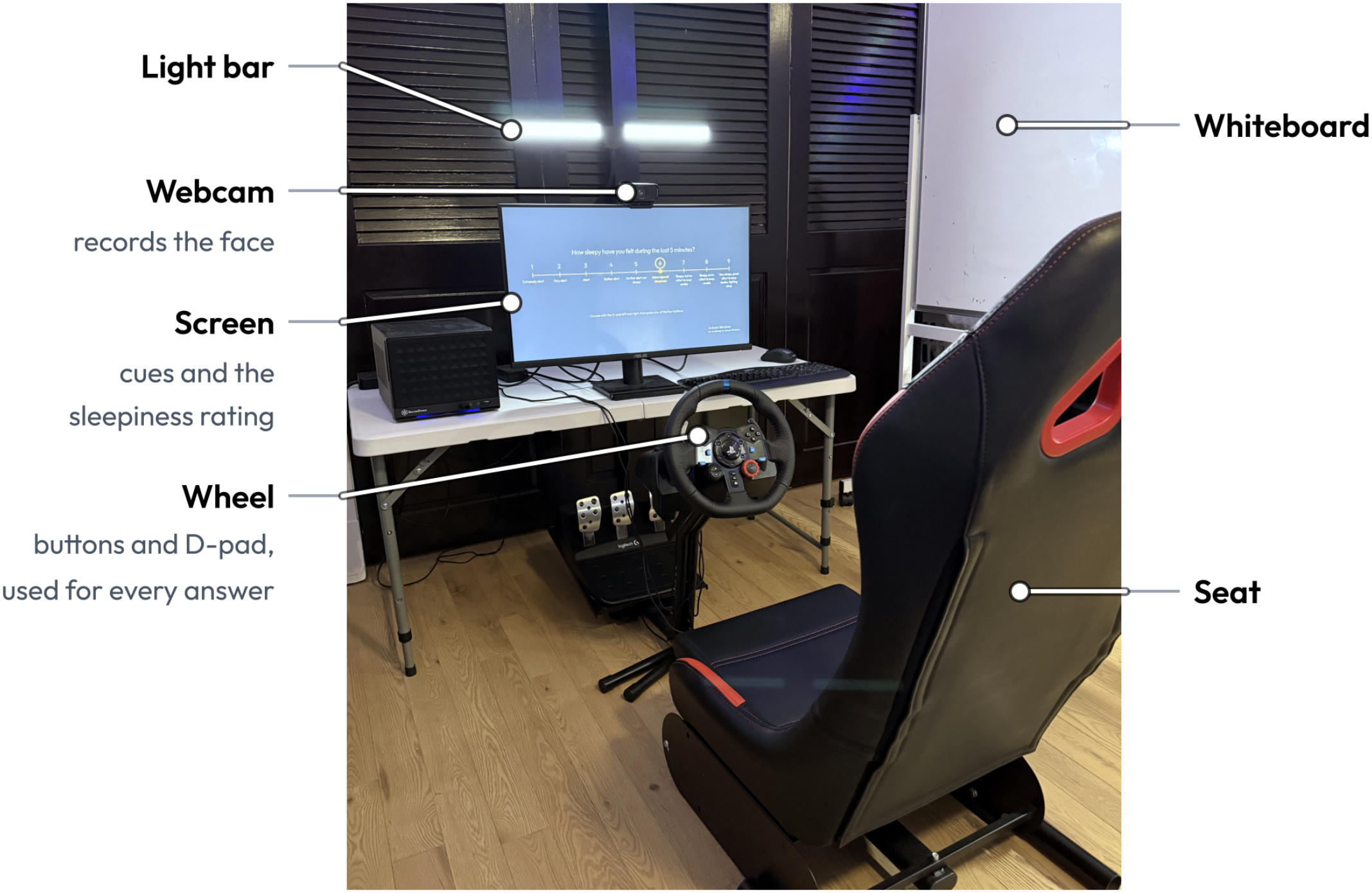
The station. The screen shows the sleepiness rating. A webcam above it records the face, with a light bar behind. Cues were answered with the four buttons on the wheel, and ratings set with its direction pad (D-pad). A whiteboard stood to the participant’s right, and a member of the study team sat behind the seat, out of frame. The pedals were not used.

An eyes-open and eyes-closed block preceded the task, in alternating segments cued by voice, and was repeated after it (Fig. 3). The block supplies each participant’s eyes-closed reference for the personal EEG detector of Section V-F and the calibration of the camera’s eyelid measures. In the block before the task, power in the alpha band (8 to 12 Hz) on the TP9 minus TP10 derivation was higher with eyes closed than with eyes open in eight of the nine participants (median ratio 1.8, range 0.76 to 6.9; Fig. 4).

**Fig. 3.**
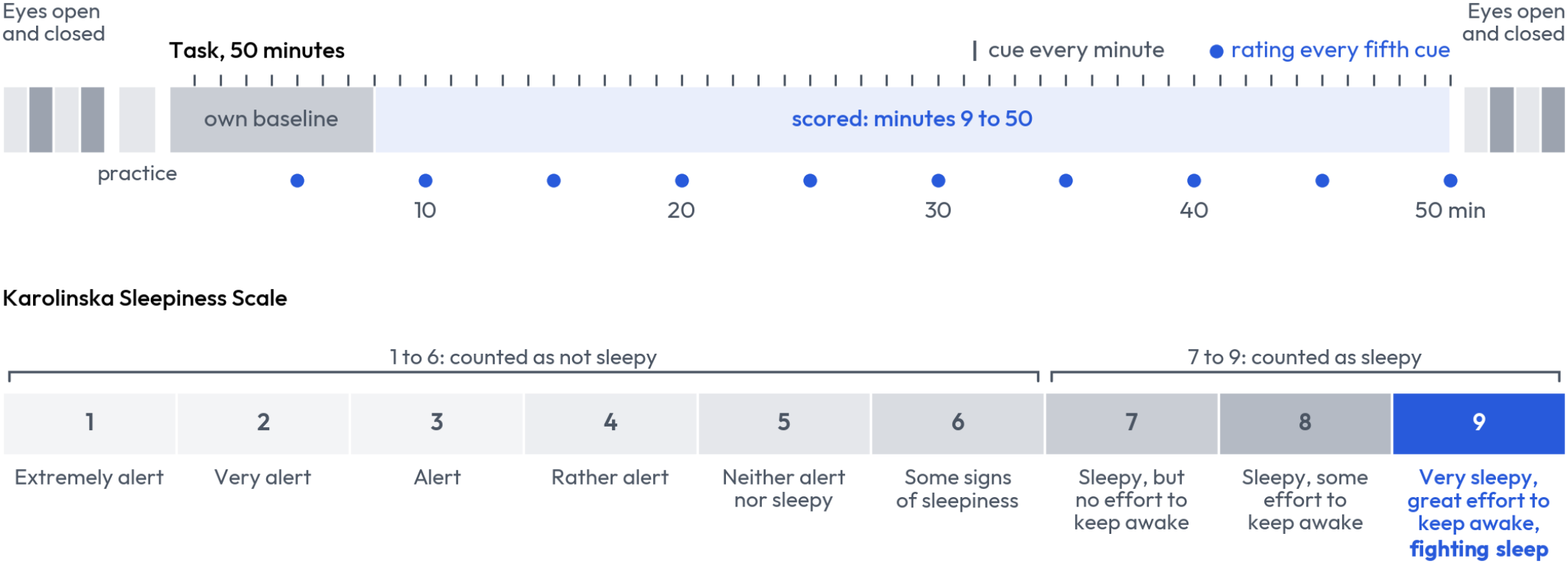
Session timeline and rating scale [15], as worded on screen. The first eight minutes of the task are each participant’s own baseline. A rating of 9 is called fighting sleep throughout, and the shading is the one used for ratings in later figures.

**Fig. 4.**
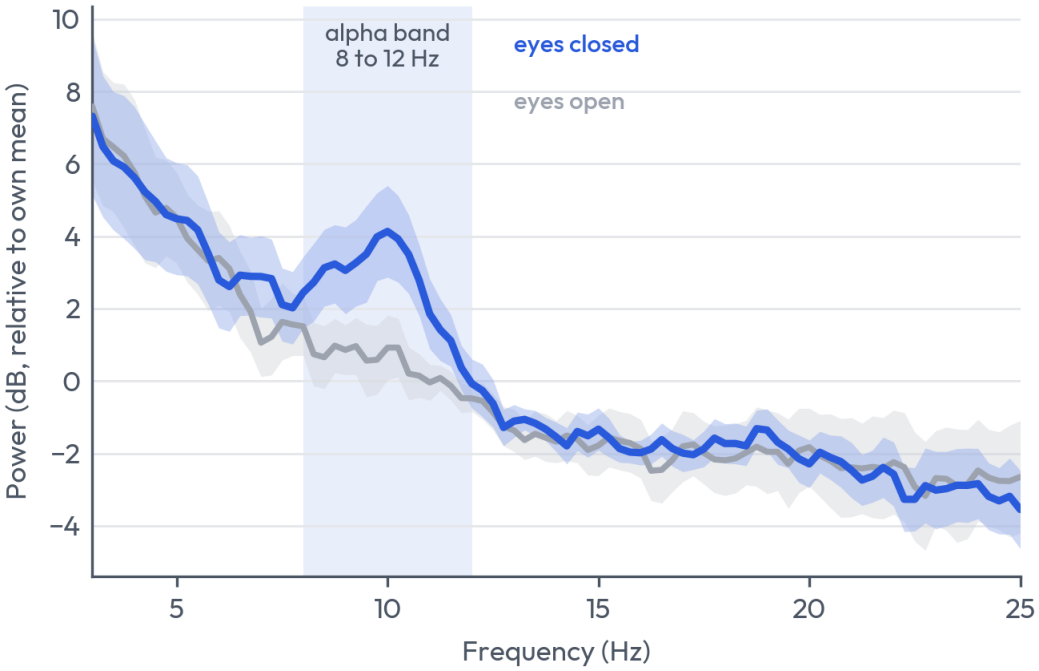
Alpha rhythm. Average spectrum at the temporal electrodes (TP9 minus TP10) with eyes closed and with eyes open, in the four minutes before the task. All nine participants (twelve recordings; a participant’s two visits are averaged; each participant’s spectra are centred on their own mean level; bands are one standard error). Power in the alpha band (8 to 12 Hz, shaded) rises when the eyes close. It was higher with eyes closed in eight of the nine participants.

Subjective sleepiness was rated on the nine-point Karolinska Sleepiness Scale [15] at the start of the session, after the practice block, every five minutes during the main block and at the end. A rating is taken to cover the minutes since the previous one. The rating selector opened on a randomly chosen value each time, so a rating could not be given by accepting a default or by repeating the last position.

A 1080p webcam recorded the participant’s face at 60 frames per second for the whole session. The video serves as a reference and a comparator only. No camera measure is an input to any headset index.

### D. Task and behavioural references

The main block is a reaction-time task. After a short practice block, a cue is presented every 60 seconds, 50 times, and reaction time is recorded for every cue. A cue with no response, or a response after more than 10 s, is counted as unanswered. The event rate is deliberately low: the state under study is produced by monotony [47], and a conventional psychomotor vigilance task with an interval of a few seconds delivers a micro-arousal at that same interval.

Fatigue is defined from behaviour alone, and no sensor data enters the definition. The behavioural fatigue score is continuous: at each cue, the mean of the within-recording rank percentile of reaction time and that of the sleepiness rating. Its most and least fatigued thirds are the two classes wherever a ranking score is reported, and the score itself is the target wherever a correlation is reported. Both components are established markers: slowed responses on a reaction-time task track sleep loss [48], and the sleepiness rating tracks both performance and physiological measures of sleepiness [19], [49]. Their combination into a single rank index is defined here and has not been validated outside this study. Stricter definitions, up to minutes rated 9 with the cue still answered, are scored in Section VI-C, and that top-of-scale definition is the one fixed for the driving trial (Section IV-A). Baseline reaction time is the mean of the first five cues. Where an onset of drowsiness is needed, it is the first cue at which reaction time exceeds 1.25 times baseline with a rating of 6 or more, on two of three consecutive cues.

### E. Recording, synchronisation and inclusion

In one recording the headband’s timestamps ran fast, and they were corrected from the device’s sample counters with no samples lost. The corrected timeline is used in Sections VI-B and VI-C. The trained detectors of Section VI-E were run on that recording before the correction, which moved their means by 0.01 to 0.02.

No recording is excluded. All twelve are reported and all twelve enter every headset analysis. Video is usable for ten recordings from seven participants; one recording’s video file was lost and one stopped early. Only minutes 9 to 50 of the main block are scored, because minutes 1 to 8 are each recording’s own baseline (Section V-B).

### F. Driving trial

The driving trial was a single outing on 17 November 2026 by one member of the study team, alone in a personal passenger car, under one uninterrupted headband fit. It had two parts. In the first the participant drove rested for 33 minutes on local streets, in daylight in the morning. In the second the car was parked with the engine idling, and the participant completed the laboratory session of Sections IV-C and IV-D in the driving seat, answering on a keyboard, with a laptop camera on the face. The headband was not removed or refitted between the parts, since refitting would confound the electrode contact with the change in setting.

The driving part tested two things: whether the headband returns a usable value for each minute in a moving car, and whether the alarm stays quiet on an alert driver. Minutes were marked every 60 s from the moment the car first moved. The first eight were the baseline and the 25 after them were scored. Blinks during the drive were detected with the threshold set in the eyes-closed minute of the parked part, since the drive has no such minute. The parked part is where sleepiness could develop, and it was scored as the laboratory sessions were.

No fatigue was induced while driving. The drive began rested, and nothing was touched or looked at while the car moved: not the headband, the computer or the task. The rule was to park at a rating above 5, or at any yawning, heavy eyelids or drifting. The participant did not drive after the parked part.

Head motion was recorded throughout by the accelerometer and gyroscope of the headband, sampled at 52 Hz. Because braking, cornering and slopes register as head tilt, the eye component of the index is reported beside the full index. The index, its alarm line and two pass marks for the drive were fixed before any driving recording existed (Section IV-A): a valid index value in at least 90 percent of driving minutes, and at most a quarter of scored driving minutes over the alarm line. That line, 0.79, is the level that 10 percent of the not-sleepy laboratory minutes go above. With one participant, the parked part is reported descriptively.

### G. Ethics, consent and data handling

No institutional review board or independent ethics committee reviewed this work. The work was done by a private company and is not affiliated with any university. The consent form said so on its first page and again in the statement each participant signed.

Each participant gave written informed consent on a form that is also the data use agreement between the participant and the company. Participants were adults with no history of seizures or neurological diagnosis and no medication that significantly affects alertness or sleep. No one was deprived of sleep. The risks were those of a long, dull task: drowsiness, and mild discomfort from the headband. A member of the study team stayed in the room throughout, participants could stop at any time, and a ride home was offered to anyone too drowsy to drive.

Some participants are members of the company. The form states that declining or withdrawing has no effect on employment, equity or standing, and no member of the study team countersigned their own form. Participants received $30, as a gift card or goods, whether or not the session completed.

Recordings are stored under a study code. The file linking names to codes is kept separately, and access is limited to the study team. The form tells participants that their data are kept indefinitely for the company’s research and development, will not be sold or released as a dataset, and will be deleted on request. It also tells them that results would be posted on a public preprint server. No video frame or photograph of a participant appears here.

## V. Analysis

### A. Per-minute measures from the headset

The unit of analysis is the minute. Each measure is computed from the 60 s before a cue, which gives one row per cue and 50 rows per recording. The measures fall into three groups.

Eye signs come from the two frontal electrodes, AF7 and AF8, which against the Fpz reference carry the electrical signal of the eyelids and eyes. They are the length of blinks, which grows with drowsiness [17], [18], [20], and the power of slow horizontal and of slow vertical eye movements, which appear as sleep approaches [15], [16]. Blinks are detected on the mean of AF7 and AF8, band-passed from 0.5 to 8 Hz, as peaks more than six times the noise level of the participant’s own eyes-closed minutes, and blink length is the median width of those peaks at half height. Slow eye movement is the power between 0.2 and 0.6 Hz, on AF7 minus AF8 for horizontal movement and on their mean for vertical. Head signs, which camera systems read as nodding [12], come from the accelerometer: the tilt of the head from its median posture over the first ten minutes of the block, as the median and as the largest value over the minute. The EEG sign is the ratio of theta plus alpha power to beta power on the TP9 minus TP10 derivation, the conventional spectral index of drowsiness [50], [51]; it is computed from the 30 s before the cue, in the epochs described below.

The trained detectors of Section V-F use a wider set of 221 features. At each of seven sites (the four electrodes, the two left-right differences and the mean of the temporal pair) the 30 s before each cue, in 4 s epochs with any epoch over 150 µV peak to peak rejected, gives measures of band power, ratios between bands, the shape of the spectrum, and how quickly and irregularly the signal varies. Five eye and five head-motion measures are added.

### B. The fixed index

The headset index involves no training. Because these measures differ widely between individuals [20], each sign is expressed as its change from the recording’s own median over minutes 1 to 8, divided by the spread of that change across the other participants, limited to plus or minus 6 and averaged over the trailing five minutes. The three eye signs are averaged, the two head signs are averaged, and the index is the mean of the two. The direction of every sign was set from the literature beforehand: longer blinks [17], more slow eye movement [16] and a greater head tilt [12] count toward fatigue. The index sees no clock and no camera measure, and nothing from the scored participant enters a constant. A version with the EEG sign as a third component is reported beside it. How the index moves with the sleepiness rating is reported in Section VI-B: its mean over the scored minutes is 0.33 at a rating of 6 and 1.20 at 9, and in four of the five participants who reached 9 it was higher at 9 than at any lower rating they gave. For any recording made after the laboratory set, the divisor of each sign is a fixed number, its value over the ten laboratory recordings with video: 57.5 ms for blink length, 0.65 and 0.82 for horizontal and vertical slow eye movement (log power), and 8.0 and 12.7 degrees for the median and the largest head tilt.

### C. Camera reference

Face landmarks were extracted from the video with MediaPipe [52], [53] on a tracked face crop at 30 frames per second, and eyelid closure was calibrated on each participant’s eyes-open and eyes-closed minutes. The camera measures scored are PERCLOS, the share of the minute with the eyes at least 80 percent closed [11], with an alarm level of 15 percent, a threshold used in published camera systems [13] (the one fielded truck system with published settings gave its full warning at 12 percent [54]); a composite of eyelid and head-pose signs built exactly as the headset index is; head pose alone; and a 1.5 s eye-closure rule, the threshold Seeing Machines publishes for its Guardian driver-monitoring system [55]. The camera is a consumer webcam and not a commercial driver-monitoring system, and the 1.5 s rule is tested here on our webcam data, not in the commercial system.

### D. Measures

Four measures are reported for every index. The ranking score is the area under the ROC curve for the most against the least fatigued third of a recording, which is the probability that the index ranks a fatigued minute above an alert one [56]. The beyond-time correlation is the rank correlation between the index and the behavioural fatigue score over the scored minutes after a linear trend in minute number has been removed from both; it is the primary measure, because in a single unbroken session fatigue and elapsed time rise together [21], [26] and any drifting quantity earns a ranking score. The forward value is the same correlation between the index and reaction time one to five minutes later. The added value is the beyond-time correlation with the other device’s indices also removed.

Each is computed per recording, averaged within participants and then across participants. An alarm is the index exceeding, for two minutes running, the level that one in ten not-sleepy minutes (rated 6 or less) pass. Where the camera is given an alarm, PERCLOS is averaged over five minutes as the index is and its line is set the same way. In Section VI-C alone the ranking score is pooled over all scored minutes of all recordings, since a strict definition of fatigue leaves too few minutes per recording. Elapsed time, as the minute number, is scored beside every index as a clock.

### E. Chance levels and intervals

Chance needs care here, because two series that both drift over a session can correlate strongly even when they are unrelated [57]. Each p-value therefore compares the real result with results from 2,000 surrogate fatigue records (1,000 for the forward value). Each surrogate has the same values and the same slow and fast fluctuation as the real record, and can drift up or down, but has no link to the headset. They are made by keeping the size of each fluctuation and scrambling its timing [58], [59], after joining each record to its time-reversed copy so that a surrogate can carry a net drift. P-values are one-sided. In Section VI-C chance is set instead by pairing each recording’s index with another participant’s fatigue record 5,000 times, which keeps every time trend and breaks only the link to the person [57]. P-values there are corrected over the definitions scored, and intervals resample participants. Elsewhere p-values are uncorrected.

### F. Trained detectors

Trained detectors were fitted leaving one participant out [27] on four signal sources, alone and in combination, with and without elapsed time: spectral EEG, eye measures, motion measures, and a personal EEG detector. The personal detector is referenced to the eyes-open and eyes-closed baseline of each participant, recorded before the task in the same session. At TP9, TP10 and TP9 minus TP10, the log power spectrum from 2 to 30 Hz is averaged over the eyes-open segments and over the eyes-closed segments, and the difference between the two gives the eyes-closed direction for that person. The spectrum of the 30 s before each cue is then scored along that direction, so that 0 is the eyes-open spectrum of that person and 1 the eyes-closed spectrum. Nothing from the task enters the baseline. A search over 2,208 pipelines, with the choice nested inside the cross-validation [28], measured how much selection alone could add.

### G. Signal amplitude at the electrodes

For the participant recorded in both settings, the signal at the electrodes is compared between the laboratory, the parked car and the moving car. Spectra are computed at each electrode over 4 s windows with half overlap, and the median over windows is taken so that a few large transients do not set the level. Three stretches of a recording are compared: the minutes with eyes open and eyes closed before the task, the first eight minutes of the task, and the drive. The drive is divided into the seconds in which the car was moving and those in which it was standing still, told apart by the vibration at the headband’s motion sensor (more than 15 mg above 2 Hz). The size of the signal is given in microvolts, as its root-mean-square amplitude between 1 and 40 Hz over 4 s stretches, called epochs below. An epoch counts as within range when the signal spans 150 µV or less from its lowest point to its highest.

## IV. Results

### A. Recordings

Table III lists the twelve recordings, and Fig. 5 shows the rating for every minute of each. Sleepiness developed in most: the peak rating was 8 or more in ten recordings, and five participants reached 9, the top of the scale, in six recordings. Two participants stopped answering for stretches, for 15 and for 8 cues. The headset yielded an index for every scored minute of all twelve recordings; the camera chain lost two.

**TABLE III.** Laboratory recordings.

| **Visit** | **KSS at start** | **KSS peak** | **No answer** | **Over 2×** | **Video** |
|---|---|---|---|---|---|
| 1, first | 4 | 7 | 3 | 1 | Yes |
| 1, second | 6 | 8 | 0 | 3 | Yes |
| 2, first | 4 | 9 | 15 | 11 | Yes |
| 2, second | 4 | 9 | 0 | 3 | Yes |
| 3 | 6 | 8 | 0 | 0 | Yes |
| 4 | 3 | 8 | 0 | 3 | Yes |
| 5, first | 5 | 8 | 0 | 0 | Yes |
| 5, second | 8 | 9 | 0 | 3 | Yes |
| 6 | 4 | 7 | 0 | 0 | Lost |
| 7 | 3 | 9 | 0 | 6 | 17 min |
| 8 | 6 | 9 | 8 | 10 | Yes |
| 9 | 8 | 9 | 0 | 4 | Yes |

*Visit: participant number and visit. KSS peak: the highest rating given during the main block. No answer: cues with no response within 10 s. Over 2×: answered cues slower than twice the mean of the first five cues, including those answered after 10 s. Each recording has 50 cues. Participant 1's three unanswered cues on the first visit fall in the last four minutes, which are not scored (Fig. 5). The counts in Section VI-C use only scored minutes, and only cues answered within 10 s as over 2×, so they are lower.*

**Fig. 5.**
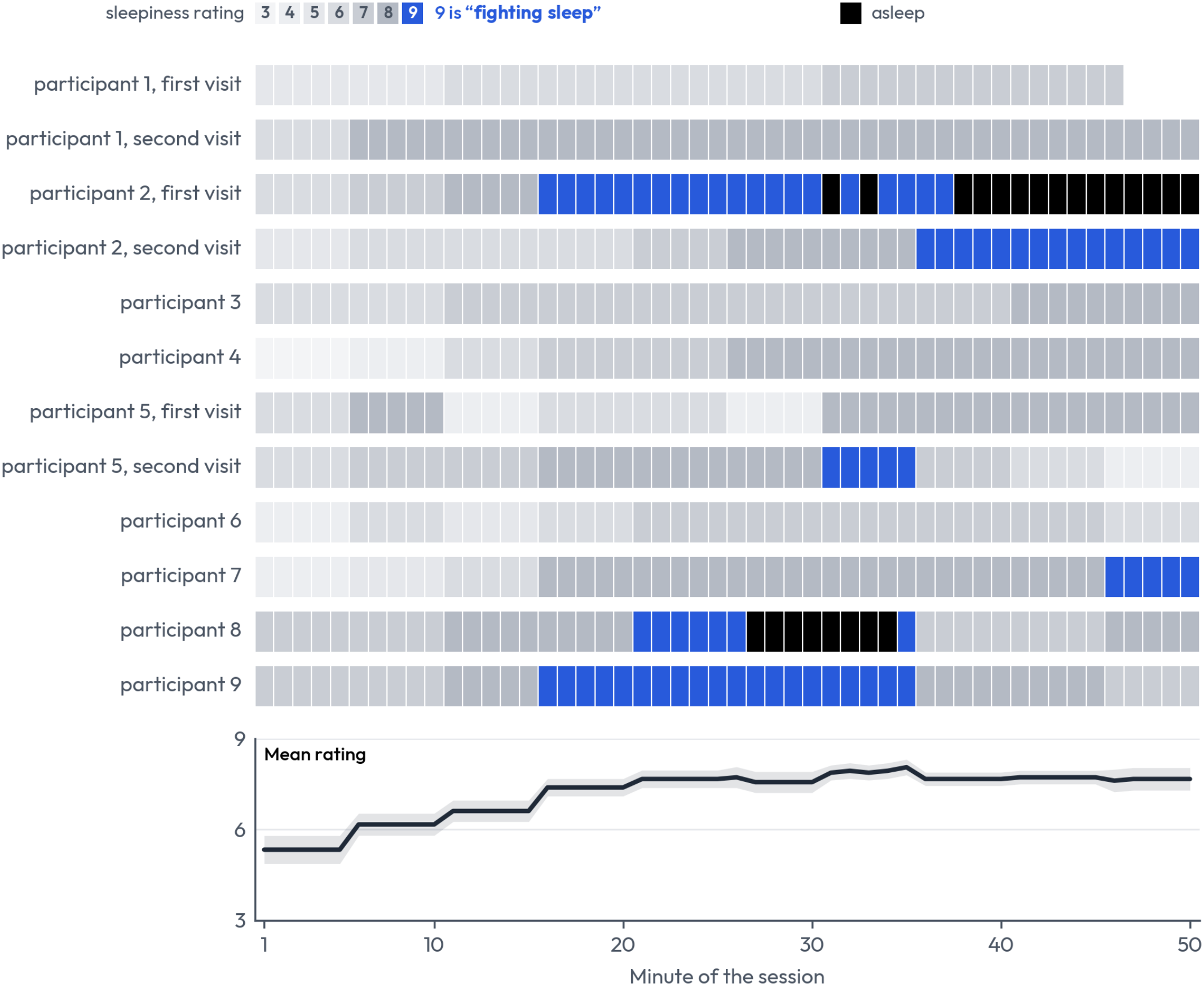
The recordings. Self-rated sleepiness for every minute of the twelve recordings from nine participants (three came twice). Blue: rated 9, fighting sleep (72 scored minutes, five participants). Black: asleep, where the cue went unanswered for 10 seconds (23 minutes, two participants). Each of these minutes was checked against the video: in the minute before the cue, the camera saw the eyes closed 42 to 98 percent of the time. Participant 1 left four minutes early on the first visit. Bottom: the mean rating by minute over the nine participants, with the two visits of a participant averaged first and minutes asleep left out; the band is one standard error.

### B. Following fatigue beyond elapsed time

People tire as a session goes on [21], [26], so a clock is a fair first guess at fatigue. Given one of a participant’s most fatigued minutes and one of their least fatigued, the minute number alone picks the right one 64 times in 100 (a ranking score of 0.64, averaged over the nine participants). The headset index picks it 73 times in 100 (0.73, p = 0.001 against chance).

The index also follows fatigue beyond what the clock explains. With a linear time trend removed from both, the index still correlated with fatigue: the beyond-time correlation is 0.32 across the nine participants (p = 0.002). It is above zero for eight of the nine and at zero for the ninth, and it stays between 0.28 and 0.36 whichever single participant is left out (Table IV). This is a result about the group and not about any one session. Second visits were mixed: two of the three sit at zero (*−*0.01 and *−*0.04) and the third at 0.47.

**TABLE IV.** Index by participant.

| **Participant** | **Recordings** | **Beyond-time correlation** | **Ranking score** | **Clock** |
|---|---|---|---|---|
| 1 | 2 | 0.27 | 0.72 | 0.74 |
| 2 | 2 | 0.25 | 1.00 | 1.00 |
| 3 | 1 | 0.40 | 0.83 | 0.54 |
| 4 | 1 | 0.23 | 0.53 | 0.86 |
| 5 | 2 | 0.56 | 0.83 | 0.55 |
| 6 | 1 | 0.16 | 0.61 | 0.63 |
| 7 | 1 | 0.00 | 0.39 | 0.69 |
| 8 | 1 | 0.59 | 0.91 | 0.43 |
| 9 | 1 | 0.40 | 0.73 | 0.36 |
| Mean over people | 12 | 0.32 | 0.73 | 0.64 |

*Ranking score: area under the ROC curve, most against least fatigued third of each recording. Clock: the same score for the minute number. A participant's recordings are averaged first.*

Fig. 6 shows one session minute by minute. Fig. 7 sets the six recordings that reached fighting sleep beside the six that did not: on the same task and the same clock, the index of the first group climbed past the alarm line and that of the second stayed below it on average.

**Fig. 6.**
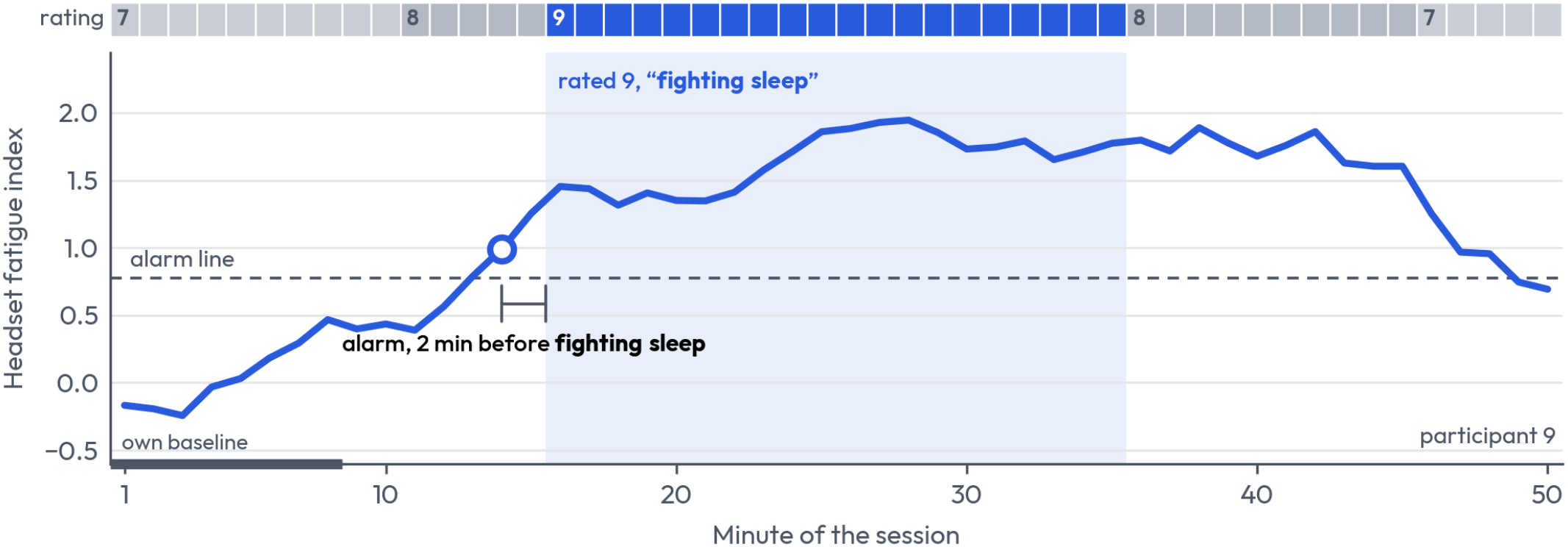
One session. Top: the sleepiness rating. Below: the headset index, as its change from the participant’s own first eight minutes. The index climbs as the rating climbs, stays high through fighting sleep and falls as the rating drops back to 8 and 7. The alarm (circle) came at minute 14, two minutes before fighting sleep began. Participant 9, whose beyond-time correlation (0.40) is the median of the twelve recordings. Dashed: the alarm line of Section V-D.

**Fig. 7.**
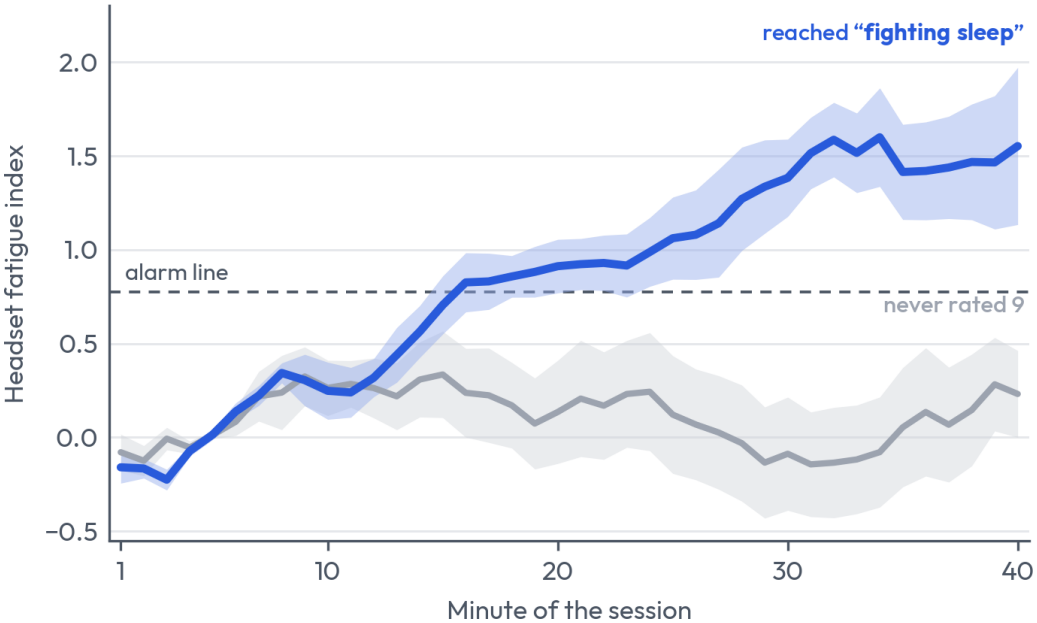
Sleepy and not-sleepy recordings. Mean headset index by minute of the session for the six recordings that reached fighting sleep and the six that never did. Bands are one standard error across recordings. All twelve sat the same task for the same time.

The eye and head measures each correlated with fatigue on their own; the EEG measure did not. The eye signs give 0.30 (p = 0.002), the head signs 0.19 (p = 0.023) and the EEG sign 0.00 (p = 0.49); adding the EEG sign to the index pulls it down to 0.23 (p = 0.013). There is also a small sign that the index runs ahead of behaviour. Its forward value, the link to reaction time one to five minutes later, is 0.06 across all nine participants (p = 0.070) and 0.09 across the seven with video (p = 0.020). The index also tracks each half of the behavioural fatigue score on its own. On the ten recordings with video, with elapsed time removed, it follows the sleepiness rating at 0.39 (p = 0.002) and reaction time at 0.25 (p = 0.006). Over all twelve recordings its average level climbs with the rating, from 0.33 at a rating of 6 to 0.50 at 7, 0.52 at 8 and 1.20 at 9.

### C. Detection as fatigue is defined more strictly

As fatigue becomes more severe, the headset index does increasingly better than the clock. Table V scores the same fixed index, with nothing fitted, against definitions of increas-ing strictness, pooled over all scored minutes. On the loose definitions it is level with a clock. At a rating of 8 or more it scores 0.70, which is not clear of chance once the number of definitions is counted (p = 0.23). At the top of the scale, minutes rated 9 with the cue still answered, it scores 0.90 (95 percent interval 0.66 to 0.97) where a clock scores 0.57, and 78 percent of those minutes lie above a line that 11 percent of all other minutes pass. Counting the minutes in which the cue went unanswered as fatigued as well lifts this slightly, to 0.91 over 95 minutes, with 80 percent detected.

**TABLE V.** Index by fatigue definition.

| **Fatigued minutes are those with** | **Minutes (people)** | **Headset** | **Clock** | **Detected** |
|---|---|---|---|---|
| Most against least fatigued third | 168 (9) | 0.66 | 0.69 | 32% |
| Sleepiness rated 7 or more | 380 (9) | 0.66 | 0.73 | 40% |
| Rated 8 or more | 261 (8) | 0.70 | 0.64 | 35% |
| A response over 2× baseline | 36 (7) | 0.74 | 0.54 | 44% |
| Rated 9, cue still answered | 72 (5) | 0.90 | 0.57 | 78% |
| Cue unanswered for 10 s or more | 23 (2) | 0.90 | 0.72 | 74% |

*Headset and Clock: area under the ROC curve over all scored minutes of all twelve recordings, fatigued minutes against every other scored minute (first row: against the least fatigued third). Detected: share of fatigued minutes above the line that 10 to 13 percent of the other minutes pass.*

The severity analysis is exploratory: the definitions in Table V were compared on the same recordings, and the top-of-scale definition was then frozen for the driving trial. Against chance set by re-pairing, 0.90 was reached in none of 5,000 draws (p < 0.001 after correction over definitions), and it is not an effect of elapsed time, since comparing only minutes at the same point of the session gives 0.91. The 72 minutes come from 16 ratings of 9 by five participants. The index picks them out clearly in three (0.92 to 0.98) and poorly in two (0.66 and 0.50), so counting each person once gives 0.81. With the divisors of the index fixed as numbers, as they are for the driving trial (Section V-B), the top-of-scale score is 0.89.

A step at the top of the scale is what earlier work leads one to expect: physiological signs of sleepiness have been reported to appear mainly at the highest ratings and to change little below them [15].

### D. Against a camera

Table VI sets the headset index beside four camera indices on the ten recordings with video, and the headset index comes out on top in every column. It has the highest ranking score, 0.79 against 0.65 to 0.76 for the camera. It follows fatigue beyond elapsed time more closely than any camera index, 0.39 against 0.15 to 0.26, and that lead is significant over three of the four (p = 0.016 to 0.030). And the two headset indices are the only ones whose forward value clears chance. Two margins are not yet secure at this sample size: the ranking score is significantly ahead of the 1.5 s closure rule only (p = 0.011), and the beyond-time lead over PERCLOS (+0.14, p = 0.088) and the forward-value margins (0.03 to 0.06) fall short of significance.

**TABLE VI.**
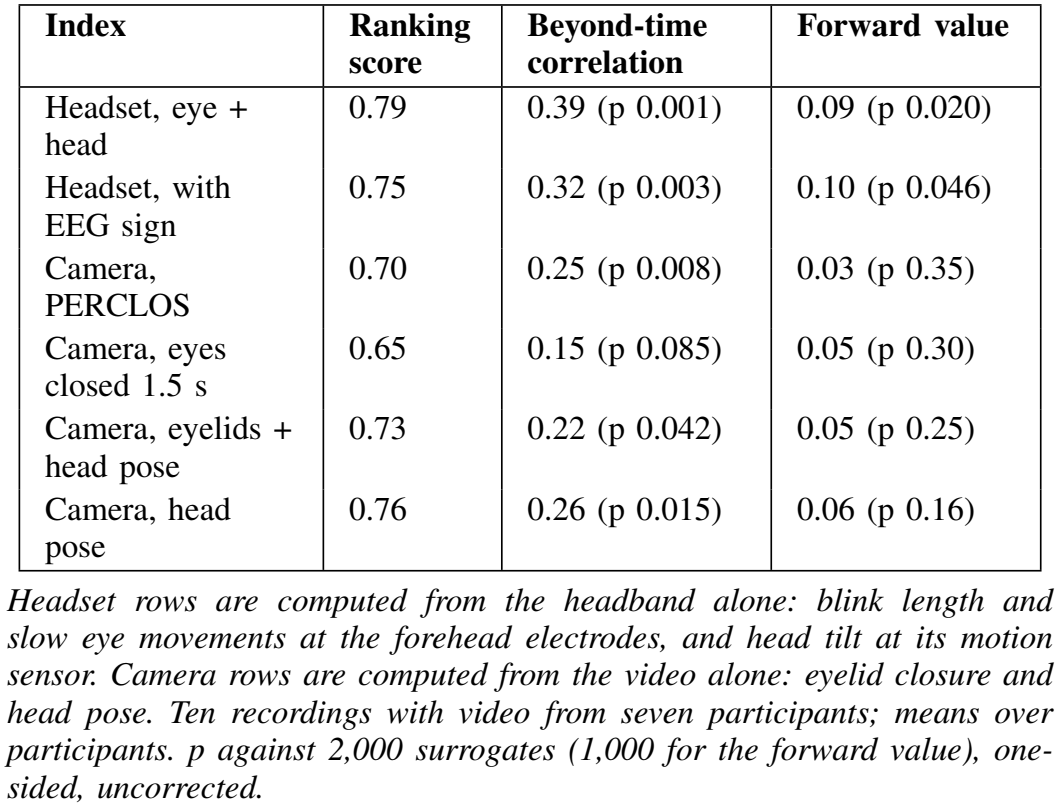
Headset and camera.

The headset carries information the camera does not. Remove everything the clock and the camera indices can tell, and the headset index still follows fatigue (0.27, p = 0.002). Remove the clock and the headset index, and the camera composite no longer clears chance (0.09, p = 0.17).

With each alarm line set for a 10 percent false-positive rate on the 73 not-sleepy minutes (rated 6 or less; 11 and 12 percent in practice), the headset index caught 91 percent of the minutes rated 9 and PERCLOS caught 33 percent (67 minutes from 15 ratings of 9 by four participants; by participant, the headset caught 97, 100, 29 and 100 percent and PERCLOS 46, 0, 43 and 15 percent; difference 58 points, 95 percent interval 6 to 89; Fig. 8). A clock, the minute number, set the same way caught 16 percent. At the line fixed for the driving trial (0.79, Section IV-F), the index with its divisors fixed caught 82 percent of all 72 minutes rated 9. At a rating of 8 or more the figures are 46 and 19 percent (interval 5 to 47), and at 7 or more 40 and 16 percent (9 to 41). On timing the answer depends on what is counted. Counted as which device alarms first in a recording for any reason, with the camera taken as the earliest of PERCLOS, the composite and head pose, the headset was first in four recordings and the camera in five, with one level. Five recordings with video reached fighting sleep. With both alarms set as above, the headset alarm had already sounded when fighting sleep began in four of them, and sounded nine minutes after in the fifth. The PERCLOS alarm sounded 6 to 13 minutes after it began in four and never in the fifth (Fig. 9). Over seconds the camera is the better instrument. Across 105 eye closures of a second or more, its eyelid measures gave warning of the closure one to five seconds ahead (0.62 to 0.63, p < 0.01), while the headset was at chance at every lead (0.43 to 0.50).

**Fig. 8.**
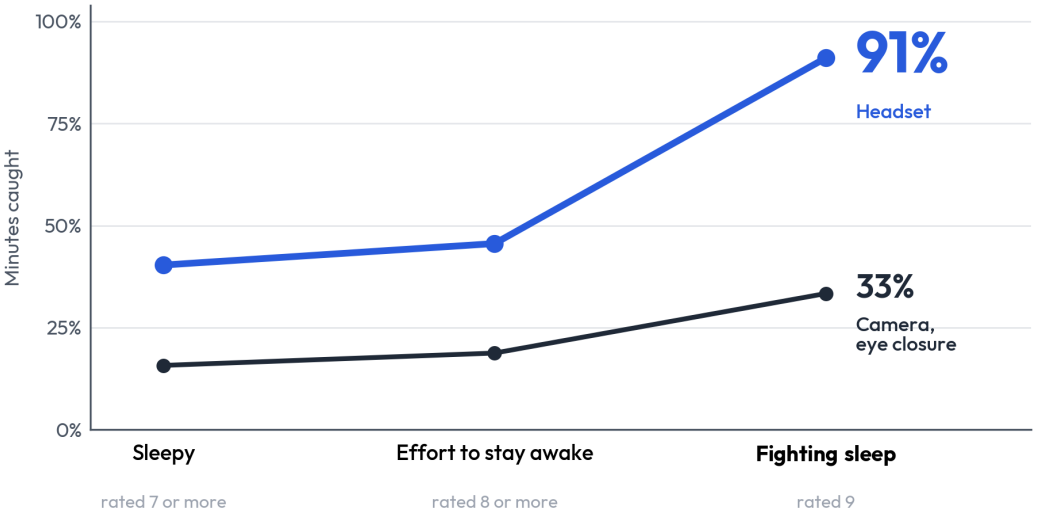
Minutes caught by sleepiness level. Share of minutes above the alarm line for the headset index and for the camera’s eye-closure measure (PERCLOS), on the same minutes of the ten recordings with video (seven participants). Each line is set so that one in ten minutes rated not sleepy (6 or less; 73 minutes) pass it. At a rating of 9 (67 minutes, four participants) the headset caught 91 percent of minutes and PERCLOS 33 percent.

**Fig. 9.**
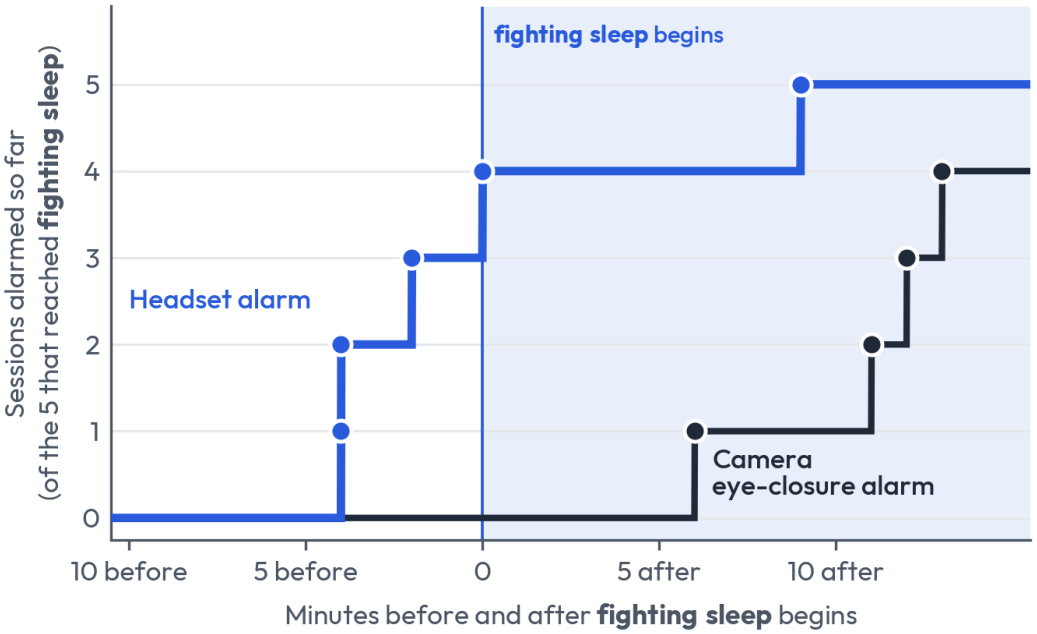
When each alarm sounded. The five recordings with video that reached fighting sleep (four participants). Each dot is the alarm of one session, placed at the minute it sounded relative to the start of fighting sleep; the line counts the sessions alarmed so far. The headset alarm had sounded by the start of fighting sleep in four of the five and nine minutes after in the fifth. The PERCLOS alarm sounded 6 to 13 minutes after in four and never in the fifth. Both are treated the same way: a five-minute average, a line that one in ten not-sleepy minutes pass, and an alarm after two minutes in a row above it.

Averaged over the six recordings that reached fighting sleep, the index was flat until about five minutes before it began and crossed the alarm line about three minutes before (Fig. 10).

**Fig. 10.**
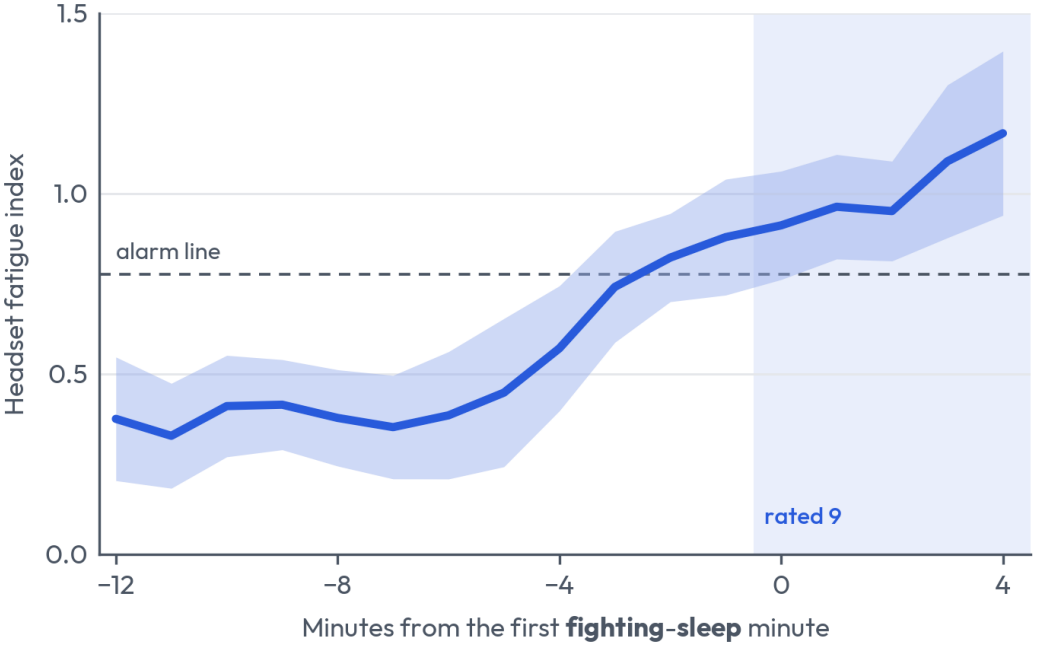
Index before fighting sleep. Mean headset index in the minutes around the first fighting-sleep minute of each recording, for the six recordings that reached a rating of 9. The band is one standard error across recordings. The index is flat until about five minutes before, then rises, and crosses the alarm line about three minutes before.

On ranking score, across the 42 definitions of Section VI-C (Table V shows six), the two devices are level. The headset index is ahead on 24 and behind on 18, and no ranking-score margin in its favour has an interval clear of zero. At the top of the scale it scores 0.93 against the camera composite’s 0.86 on the recordings with video (difference +0.07, interval *−* 0.13 to +0.23). On unanswered cues the camera is ahead, 0.98 against 0.89. On the three pairs of first and second visits, telling the more tired visit from the other, the headset index scored 0.49, 0.72 and 0.27 and the camera composite 0.50, 0.70 and 0.92.

The camera also reads one participant, who looks down between cues, as having closed eyes: 36 percent of that participant’s alert minutes are over the camera’s alarm line against 7 percent for the headset.

### E. Spectral EEG and trained detectors

The personal EEG detector is the one positive result for brain-rhythm EEG, and it weakened as recordings were added. On the first seven participants it ranked fatigue defined by reaction time alone at 0.73 (p = 0.001). Over all nine it stands at 0.65 (p = 0.009), and at 0.62 (p = 0.041) on the behavioural fatigue score. On the five recordings made after that first analysis it was at chance in four and scored 0.88 in the fifth, a figure that falls to 0.62 when that participant’s unanswered cues are left out. Its contribution to a model that already has the eye and motion measures is +0.03 (p = 0.10). Generic spectral features fared no better: a sweep of the 221 features against eight targets, run on the first five participants, found no association that survived correction for multiple testing. No trained detector beats the untrained index. Leaving one participant out, eye and motion measures score 0.57 to 0.59 and the full model without a clock 0.60 to 0.62, neither clear of chance; with elapsed time added the model reaches 0.70 where elapsed time alone gives 0.64. In the nested search the pipeline chosen inside each fold scored 0.58 (p = 0.13) where the untrained index scores 0.72, and models fitted to a participant’s own session were at chance. With nine participants there is not enough data to learn a detector that transfers between people, and a fixed index with each participant’s own baseline is the most effective analysis found.

### F. Driving trial

The index stayed quiet through the drive (Fig. 11). Two pass marks had been set before the drive. The first was a valid index value in at least 90 percent of driving minutes, and the index had one in all 33. The second was that no more than a quarter of the scored minutes be above the alarm line, and none of the 25 was: the highest value was *−* 0.02 against a line of 0.79, and no alarm sounded. The eye component alone was below its own line in every scored minute as well. Moving the minute marks 30 s either way, or halving or doubling the blink threshold, left no minute above the line. The driver reported afterwards that he never rated himself above 5 on the nine-point scale during the drive, so all 25 scored minutes count as not sleepy under every definition used here.

**Fig. 11.**
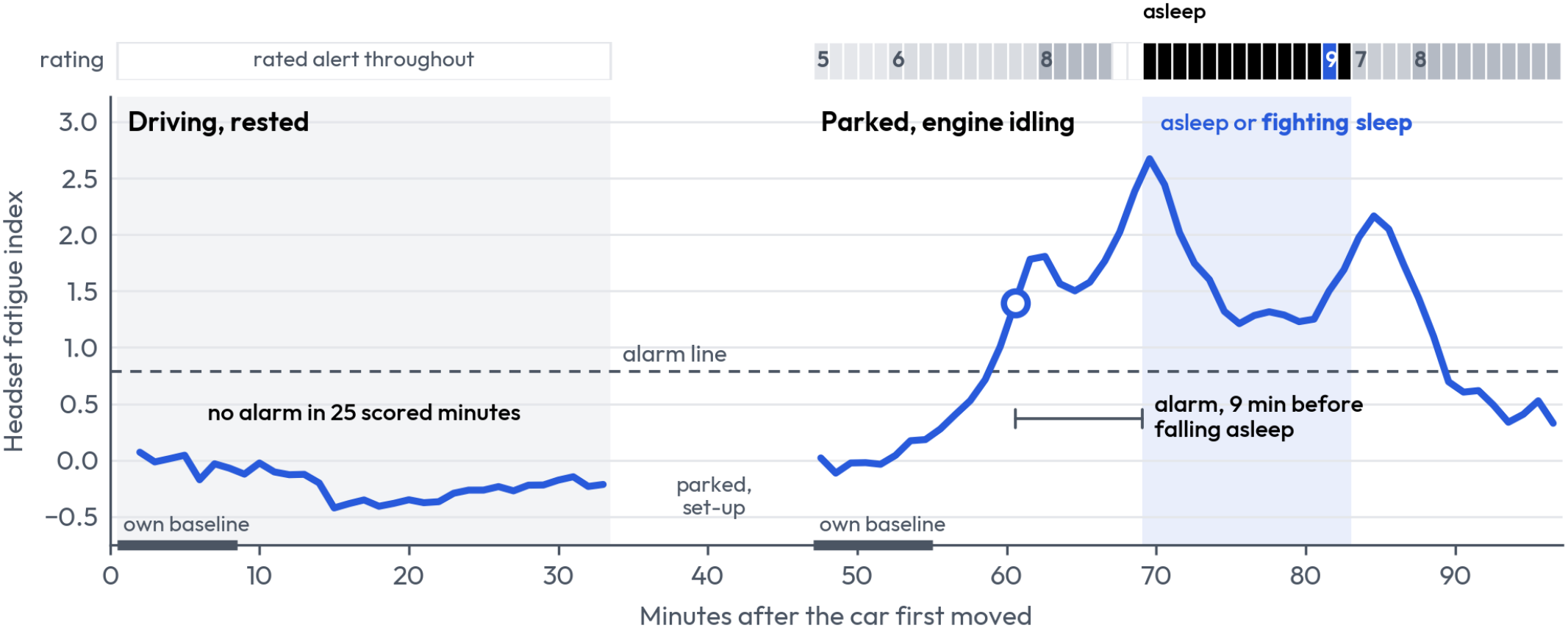
Driving trial. Left: the rested drive. Right: the same participant parked with the engine idling, under the same headband fit. Top: the sleepiness rating. During the drive the driver kept his ratings in his head, and he confirmed afterwards that he was alert throughout, never above 5. Below: the headset index, as its change from the first eight minutes of each part. The first driving minute, in which the car pulled out, belongs to the baseline and is not drawn. Circle: the alarm, at minute 14 of the parked session. Black: minutes in which the cue went unanswered and the camera found the eyes closed for 86 to 100 percent of the minute. Shaded: minutes 23 to 36 of the parked session, in which he was asleep or rated himself as fighting sleep. Dashed: the alarm line fixed for the trial. One participant, a member of the study team.

Parked outdoors with the engine idling, the participant grew sleepy and fell asleep (Fig. 12). His ratings ran 5, 6, 6 and 8 over the first twenty minutes, the next two were not given, and the one after was 9. Thirteen cues went unanswered, twelve of them in a row from minute 23 to minute 34, and in each of those twelve minutes the camera found his eyes closed for 86 to 100 percent of the time. The headset alarm sounded at minute 14 and the camera’s alarm, set as in Section VI-D with the laboratory webcam’s line applied to the laptop camera, at minute 22. The headset was eight minutes ahead. Measured against the first unanswered cue, at minute 23, the camera’s warning came with one minute to spare and the headset’s with nine. The headset alarm was also 7 minutes ahead of the first answer slower than 3 s. The index was above the line throughout the twelve minutes asleep.

**Fig. 12.**
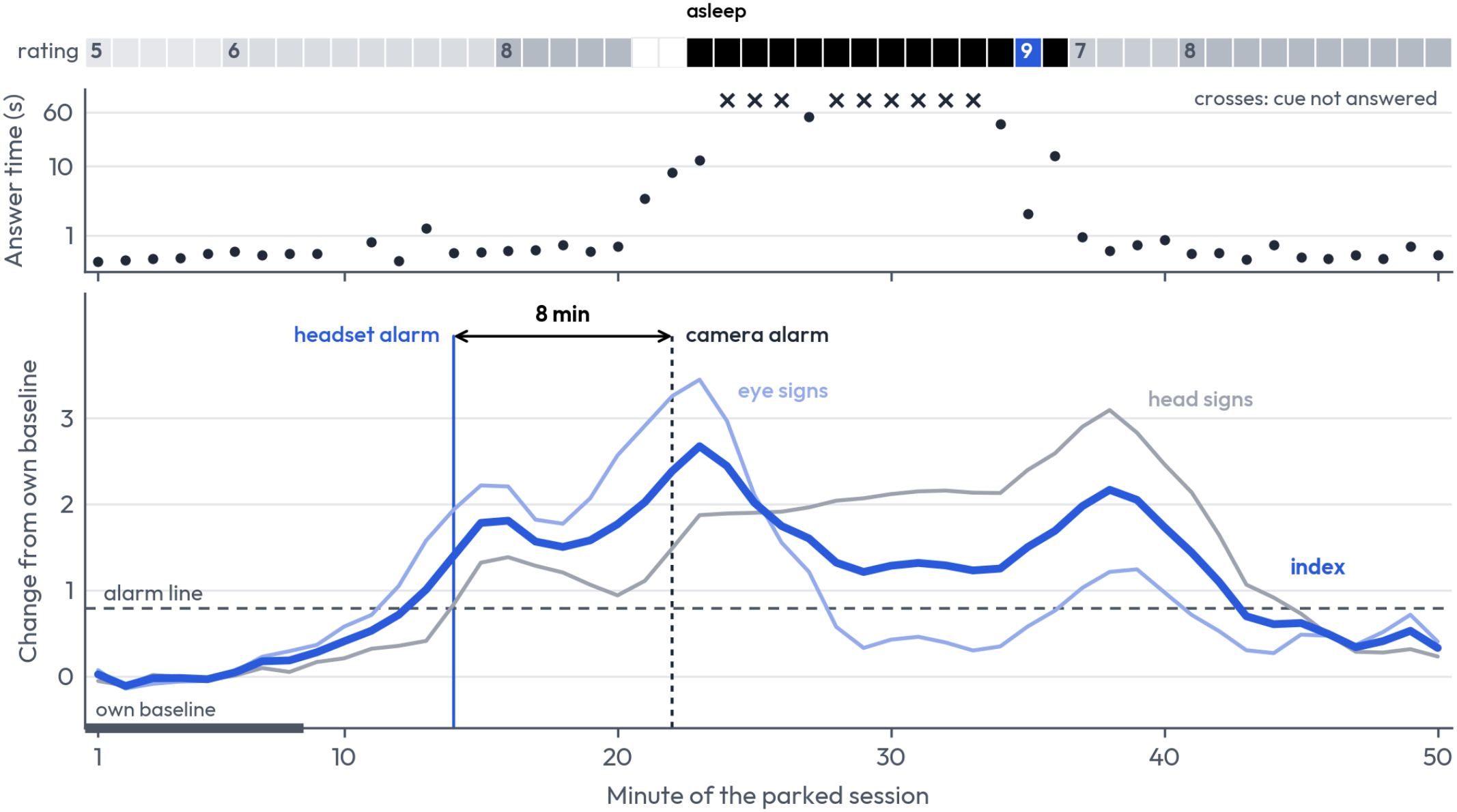
The parked session. Top: the sleepiness rating, and in black the minutes in which the cue went unanswered. Middle: the time taken to answer each cue; crosses are cues never answered. Bottom: the headset index and its two halves, the eye signs and the head signs, as change from the first eight minutes. The headset alarm sounded at minute 14 and the camera’s, set as in Section VI-D, at minute 22; the arrow marks the eight minutes between them. The eye signs rise first and fall once the eyes are shut; the head signs carry the index through the minutes asleep.

Three things qualify this. The headset alarm came while the rating was still 6: three of the seven scored minutes rated 6 or less were above the line, the three before the rating rose to 8, and the camera was above its line in none of them. PERCLOS in a single minute first passed the 15 percent level used in published camera systems at minute 12, two minutes before the headset alarm. And the eye component alone was below its line in all five minutes rated 9: with the eyes shut there are no blinks and little eye movement to measure, and the head signs carried the index through those minutes.

The quiet alarm of the drive depends on where the baseline is taken, a point examined only after the recording. Driving itself changed the signs behind the index (Fig. 13). In the first eight minutes of the drive, slow horizontal eye movement was about two of the index’s units higher than in the first eight minutes parked or in the laboratory, as scanning a road would produce. Blink length was little changed (154 ms against 142 to 147 ms), though there was no video to check blink detection during the drive. Head tilt was higher as well over those eight minutes (a median of 5.1° against 1.7° to 3.0°). The baseline fixed beforehand was the first eight driving minutes, and against it the index did not move. Against a baseline taken parked, which is not the rule that was fixed, the eye component would have been above its line in 24 of the 33 driving minutes.

**Fig. 13.**
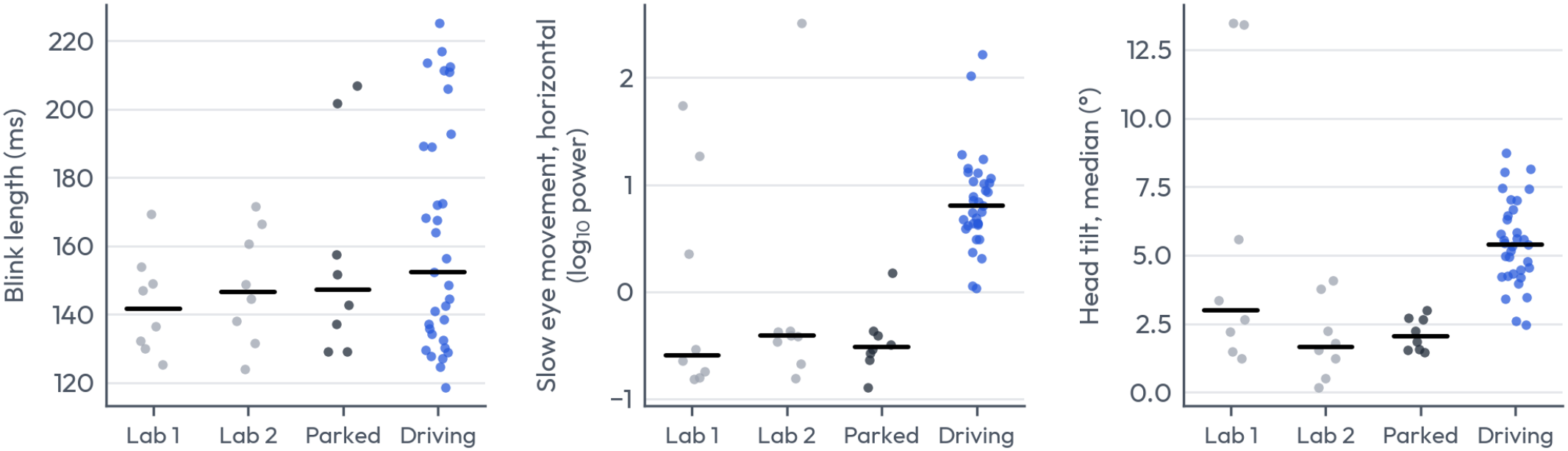
Index signs by setting. Blink length, slow horizontal eye movement and head tilt, one dot a minute, for the participant who drove: the first eight minutes of his two laboratory sessions (Lab 1, Lab 2) and of the parked session, and all 33 minutes of the drive. Bars are medians. Slow eye movement and head tilt sit higher throughout the drive; blink length changes little.

The signal at the electrodes was compared for this participant between the laboratory, the parked car and the moving car (Table VII, Fig. 14). Parked with the engine idling, the signal was no larger than in the laboratory. During the task, power at each electrode in each band from 1 to 45 Hz was 0.4 to 1.2 times its laboratory level, where this participant’s two laboratory visits differed from each other by 0.7 to 1.7 times, and the 60 Hz mains hum of the laboratory was absent. The alpha rhythm answered to closing the eyes as it had in the laboratory (Fig. 14a): alpha power at the temporal electrodes was 1.8 times higher with eyes closed before the parked task and 1.6 times after it, against 1.5 and 2.4 before the task in the two laboratory visits.

**TABLE VII.** Signal amplitude by setting.

| **Setting** | **Forehead (µV)** | **Temporal (µV)** | **Epochs in range, forehead / temporal** |
|---|---|---|---|
| Laboratory, task (two visits) | 2.6 | 5.2 | 99% / 88% |
| Parked, engine idling, task | 2.3 | 4.7 | 100% / 97% |
| Driving, car standing | 2.4 | 5.6 | 100% / 98% |
| Driving, car moving | 5.0 | 12.5 | 99% / 83% |

**Fig. 14.**
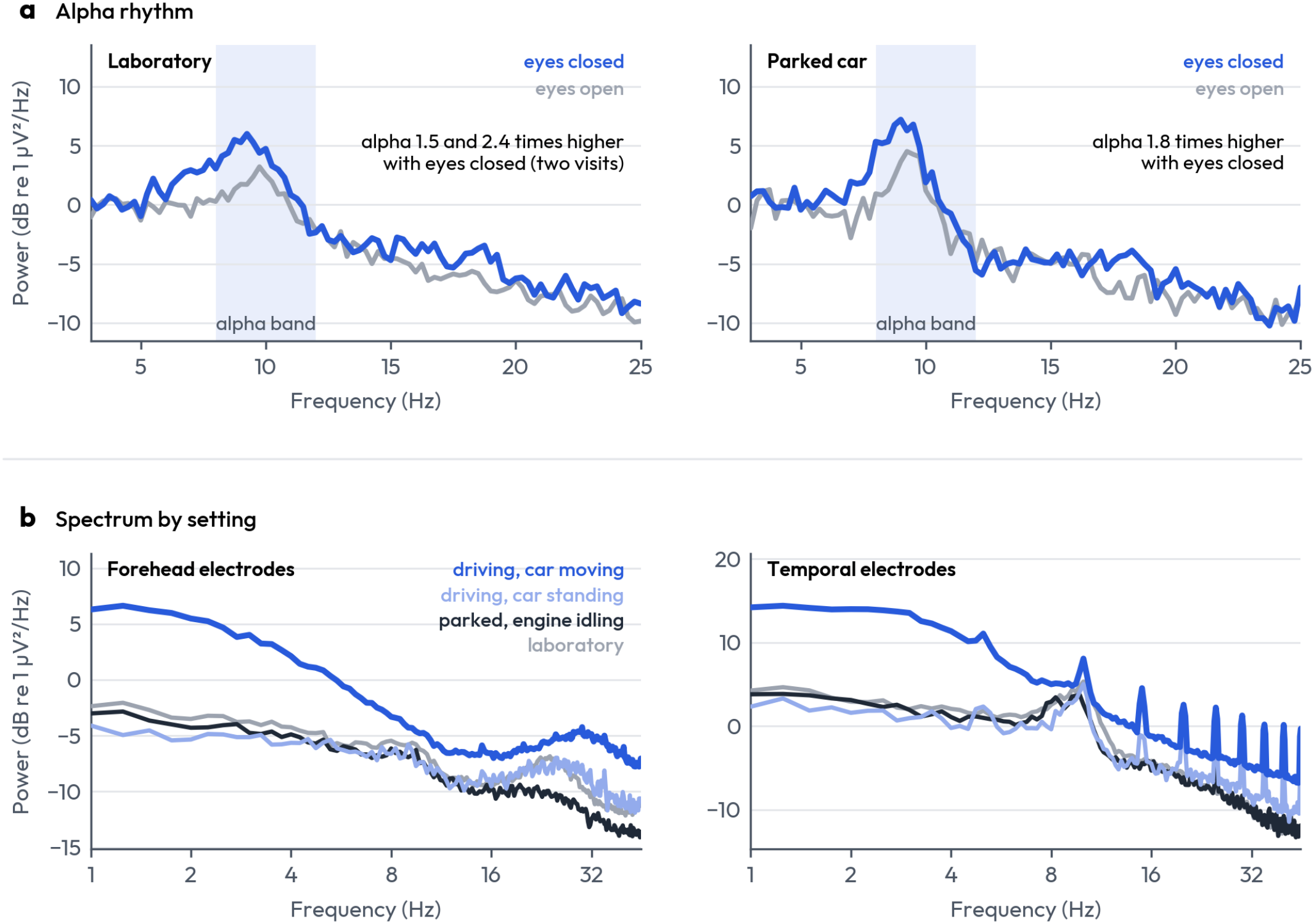
Alpha rhythm and spectrum by setting, for the participant recorded in both. a: the spectrum at the temporal electrodes (TP9 minus TP10) with eyes closed and with eyes open, before the task, in the laboratory (two visits averaged) and in the parked car; the alpha band is shaded. b: the spectrum at the forehead and at the temporal electrodes during the first eight minutes of the task in the laboratory and in the parked car, and during the drive with the car standing and with it moving. Spectra are medians over 4 s windows; in the car, over all the windows in which it was moving or standing. The narrow peaks at the temporal electrodes while the car moved are the interference at multiples of 5 Hz described in the text.

Amplitude: root mean square of the 1 to 40 Hz signal, median over 4 s epochs, mean of the two electrodes of each pair. In range: peak to peak within 150 µV.

While the car moved, amplitude about doubled, from 2.6 to 5.0 µV at the forehead electrodes and from 5.2 to 12.5 µV at the temporal ones. The added power sat mostly below 8 Hz (5 to 13 times the laboratory level at 1 to 4 Hz) and above 30 Hz (3 to 5 times), and least in the alpha band (1.1 to 1.8 times). It followed the motion of the car: in the stretches in which the car stood still during the drive, 228 s in all, amplitude was back at the laboratory level. Some of the added low-frequency power at the forehead is the eye movement of driving and not noise. The forehead electrodes, which carry the eye signs, stayed within range in 99 percent of 4 s epochs while the car moved, and the temporal electrodes in 83 percent. In the first 90 s of the recording and the last 17 minutes of the drive the temporal electrodes also carried a narrow interference at multiples of 5 Hz (the sharp peaks in Fig. 14b), about 4 µV at TP10, which was absent at the forehead electrodes and in the laboratory. It began and ended abruptly and did not follow the motion of the car, so it is electrical pick-up and not vibration; its source was not identified.

What this trial does not show is that the index detects fatigue on the road. For safety, no fatigue was allowed while driving, the one participant is a member of the study team in his third session and knew what the trial was for, and whether the alpha rhythm can be read in a moving car was not tested, since the eyes cannot be closed while driving.

## VII. Discussion

The headband caught the minutes in which people were fighting sleep. With its alarm line set for a 10 percent false-positive rate on not-sleepy minutes, the headset index caught 91 percent of the minutes participants rated at the top of the sleepiness scale. The standard eye-closure measure from the camera, set the same way on the same minutes, caught 33 percent (67 minutes from four participants; difference 58 points, 95 percent interval 6 to 89). The gap is there at lower ratings too: 46 against 19 percent at a rating of 8 or more, the level by which the Euro NCAP driver-monitoring protocol requires a driver to be classed as drowsy [14], and 40 against 16 percent at 7 or more.

The index did not simply follow time on task. All twelve recordings sat the same task for the same fifty minutes. In the six that reached fighting sleep the index climbed past the alarm line and kept climbing; in the six that never did, it stayed below the line on average. Across the whole cohort the index followed fatigue from minute to minute, beyond what time on task explains, above zero in eight of the nine participants and at zero in the ninth. It also returned a value for every minute of every session.

The index rose before the first rating of 9. Averaged over the six recordings that reached fighting sleep, it was flat until about five minutes before the first minute rated 9, then rose and crossed the alarm line about three minutes before; individual headset alarms came between four minutes before and nine minutes after. In the five such sessions with video, the headset alarm had sounded by the start of fighting sleep in four; the camera alarm, set to the same false-alarm rate, sounded 6 to 13 minutes after it began in four and never in the fifth. Counted instead as which device alarms first in a recording for any reason, the two are even (Section VI-D).

The signal comes from the eyes and the head. The index is built from blink length and slow eye movements picked up by the forehead electrodes and from head tilt measured by the motion sensor. The temporal electrodes record the alpha rhythm, which rose with eyes closed in eight of the nine participants, but brain rhythm did not add to the index on these recordings. Three limits apply. They rest on nine people and on six recordings that reached fighting sleep. The camera here is a webcam scored with the standard eye-closure measure; timed by the eye-closure cutoffs that camera vendors publish, a camera alarms as early as the headset does, but those cutoffs also sound in 26 to 53 percent of not-sleepy minutes against 11 for the headset. And the headset alarm also sounded in three of the six recordings that never reached fighting sleep.

The headband kept working in a car. On one outing by one participant it returned a value for every minute of a 33-minute drive and raised no alarm on a rested driver. With the car parked its alarm sounded nine minutes before he fell asleep and eight minutes before the camera’s. In that session a warning from the camera would have come with one minute to spare, and a warning from the headband with nine. The alarm came while he still rated himself 6; by minute 20 he rated 8, and three minutes later he stopped answering. The recording itself held up as well. With the engine idling the signal was no larger than in the laboratory, and the alpha rhythm answered to closing the eyes as before. With the car moving the amplitude doubled, but the forehead electrodes that carry the index stayed in range in 99 percent of epochs. This shows that the measurement can be made in a moving car. It does not show that the index detects fatigue there. That was left untested for safety: no one was to drive while sleepy, so the drive was made rested and the participant was parked before he could grow sleepy.

## Limitations

Nine participants took part, some of them members of the study team, and the top-of-scale result rests on 72 minutes from five people. That is a small sample, though in line with comparable studies of sleepiness and driver fatigue, which have drawn their conclusions from eight [15], nine [25], ten [16], [20], [22], twelve [60] and fourteen [23] participants. Most of those studies, however, recorded each person on two or more occasions [16], [20], [22], [23], [25]; here three of the nine came back for a second session. The analysis was exploratory and was developed on the data it reports. Each session is a single unbroken block of about fifty minutes, so fatigue and elapsed time rise together and can be separated only statistically; a session with a break in the trend would separate them by design. The index measures change from the first eight minutes and so assumes the participant begins alert, which two recordings, which began at a rating of 8, did not. The reference is a self-rating and a reaction time, not a physiological standard, although the rating has been validated against both performance and EEG [19], [49]. The camera comparator is a webcam with open-source landmarks, not a commercial driver-monitoring system. And a fifty-minute session at a simulator station says nothing yet about a ten-hour shift on the road, where sleepiness in a car has been found to run lower than in a simulator [22]. The driving trial is one outing by one member of the study team, with no rating recorded during the drive. Its quiet alarm rests on a baseline taken while driving: driving raised the slow eye movement sign by itself, and against a baseline taken parked the eye component would have alarmed in 24 of the 33 driving minutes.

## VIII. Conclusion

A dry-electrode headband worn for fifty minutes of a monotonous task yields, with no training and no clock, an index that follows fatigue beyond elapsed time across nine people and that, counted per person, ranks a minute rated as fighting sleep above another minute about eight times in ten. The signal comes from the eyes and the head, not from brain rhythm. With its alarm line set for a 10 percent false-positive rate on not-sleepy minutes, it caught 91 percent of the minutes people rated as fighting sleep, where the standard eye-closure measure from a webcam, set the same way, caught 33 percent, and its alarm had sounded by the start of fighting sleep in four of the five sessions that could be compared. It adds to the camera and it does not lose the wearer. On one outing in a car, with the same rule unchanged, the alarm stayed quiet through a 33-minute rested drive and, once the car was parked, sounded nine minutes before the driver fell asleep and eight minutes before the camera’s alarm. That is one participant, and the test on new people is still to come.

## Competing Interests

R.S.B., A.B., L.R.W. and W.M.B. are co-founders of Neuro Safety Systems, Inc., hold equity in it and are paid by it. The company is developing a commercial driver fatigue monitor.

## Funding

This work was funded by Neuro Safety Systems, Inc. The company designed and ran the study, analysed the data and wrote the paper.

## Data and Code Availability

The recordings are not available. The consent form promised participants that their data would not be deposited in a repository or released as a dataset. The analysis code is not public. The construction of the index and the alarm rule are described in Sections V-B and V-D.

